# Extracting Evidence Fragments for Distant Supervision of Molecular Interactions

**DOI:** 10.1101/192856

**Authors:** Gully A Burns, Pradeep Dasigi, Eduard H. Hovy

**Affiliations:** USC Information Sciences Institute, Marina del Rey, CA 90292, USA; Language Technologies Institute – Carnegie Mellon University, Pittsburgh, PA 15213, USA

**Keywords:** Machine Reading, Molecular Interactions, Biomedical In-formatics, Discourse Analysis

## Abstract

We describe a methodology for automatically extracting ‘evidence fragments’ from a set of biomedical experimental research articles. These fragments provide the primary description of evidence that is presented in the papers’ figures. They elucidate the goals, methods, results and interpretations of experiments that support the original scientific contributions the study being reported. Within this paper, we describe our methodology and showcase an example data set based on the European Bioinformatics Institute’s INTACT database (http://www.ebi.ac.uk/intact/). Using figure codes as anchors, we linked evidence fragments to INTACT data records as an example of *distant supervision* so that we could use INTACT’s preexisting, manually-curated structured interaction data to act as a gold standard for machine reading experiments. We report preliminary baseline event extraction measures from this collection based on a publicly available, machine reading system (REACH). We use semantic web standards for our data and provide open access to all source code.

## 1 Introduction

The biomedical literature consists of tens of millions of published articles [1] and there are thousands of informatics systems that catalog both published and unpublished scientific work [2]. These databases are typically constructed manually and there is therefore a very strong need to automate extraction of information from research articles using machine reading approaches. We are attempting to explore whether extracting and representing primary experimental evidence will provide a more accurate, and scoped target for machine reading than simply attempting to read all text in the body of a paper article with equal priority [3]. This report provides the starting point of our investigation by identifying which fragments of an experimental article’s narrative specifically describe the experimental contribution of that article.

In order to develop machine reading systems, we require training data that links the text of research papers to structured semantic representations of the knowledge content. We describe a general method for creating annotated corpora based on *distant supervision* to create links between text describing research evidence to previously-curated database records. We seek to use figure references in the text of articles to create a useful link between text and data (Figure 1).

**Figure 1.**
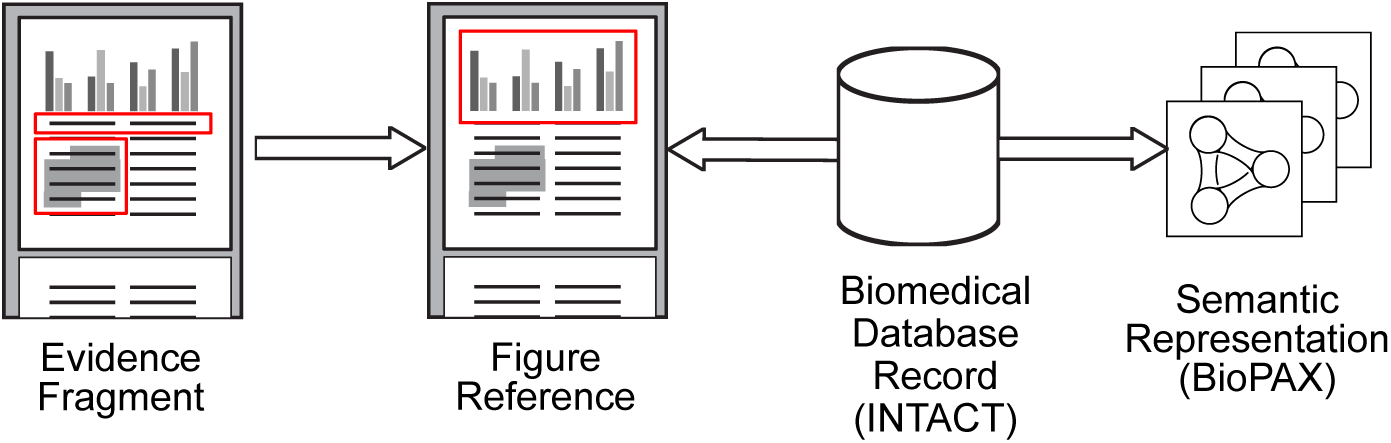
Figure references can link relevant fragments from full-text primary research articles to database records and derived semantic representations.

The European Bioinformatics Institute’s (EBI) INTACT database describes molecular interactions (binding events where two molecules join to form a complex). INTACT links each figure reference (*i.e.*, 1a, 2b, 5f, *etc.*) directly to database records [4]. Figure 1 illustrates how evidence fragments might then be linked to database records via their common figure reference.

We automated this linkage between database records and evidence fragments to provide a cost-effective way of creating corpora. We applied an open-source event extraction method for signaling pathway events (REACH) [5] to develop a baseline for detailed semantic extraction of this text.

## 2 Related Work

In biomedicine, distant supervision was originally used to facilitate entity and relation extraction from text using structured data [6]. Previous efforts center around record linkage between domain-specific biomedical entities (such as proteins and residues, see [7]). The method we use to tag discourse elements is simpler than general discourse parsing methods (such as Rhetorical Structure Theory (RST) [8]), which might be applied to open domain text. More precisely, our work mirrors that of Teufel *et al.* concerned with “Argumentative Zoning” where classifiers act on sentences across the entire narrative scope of a paper [9].We seek a more restricted focus in order to isolate a paper’s primary experimental contribution for subsequent extraction, Aydin *et al.* describes a closely-related study in which they classify passages with experimental methods with PSI25-MI terms (the same terminology used in INTACT) [10]. They focus on methodological text and the size of their annotated corpus (30 papers) reflects the important role of annotated corpora in information extraction. We suggest that our use of distant supervision could increase the size of their working corpus.

## 3 Methods

### 3.1 INTACT Data and Text Preprocessing

We only used INTACT papers that had been designated as part of the open access subset of Pubmed Central’s online digital collection. Our INTACT data contains 13,991 papers of which 1,063 were available for use. To split sentences into their constituent clauses, we computed dependency parses with the Stanford Lexicalized Parser. INTACT data was downloaded and cross referenced to the open access publications with figure references to yield 899 papers containing 6320 individual reported reactions of molecular interactions.

### 3.2 Science Discourse Tagger Neural Net Classifier

We used the *Science Discourse Tagger* (SciDT) [11] to annotate individual sub-sentence clauses from scientific papers with one of eight discourse tags including ‘fact’, ‘’problem’, ‘hypothesis’, ‘goal’, ‘’method’, ‘’result’, and ‘none’ [12]. Training data was manually compiled from 20 papers. We ran release v0.0.2 from the SciDT and SciDT Pipeline github repositories.

### 3.3 Linking Figure References to Surrounding Text

We used a rule-based approach to locate the sentence boundaries of text pertaining to specific subfigures. Figure 2 shows an example from [13]. This shows the delineation of text passages pertaining to the evidence presented in subfigures 1A, 1B and the first sentence of the description of 1C. Color coding of sentences shows the discourse tags associated with each clause shown.

**Figure 2.**
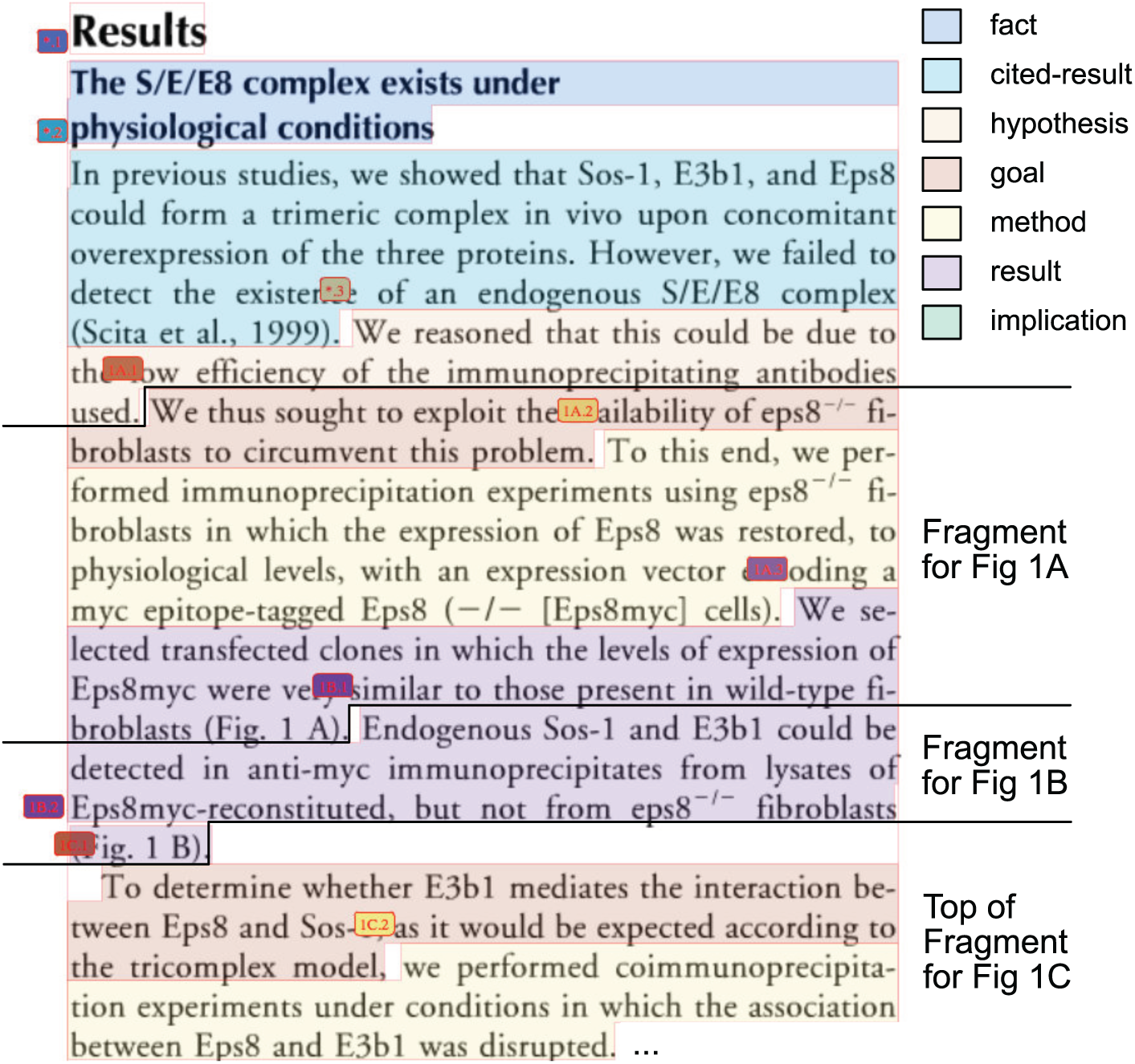
Evidence text fragments referring to subfigures 1A, 1B and 1C of [13].

Informally, the algorithm to extract these fragments is as follows:

For each subfigure reference in the text, we first scan backwards from clause containing a figure reference mention (*e.g.*, ‘Fig. 1 A’) for the start of the evidence fragment. We assert the presence of a fragment start boundary between consecutive sentences *S*_1_ and *S*_2_ (i.e., *S*_2_ is the first sentence of the evidence fragment) if the following conditions are met:

Sentence *S*_1_ contains either

a. clauses that are tagged as ‘hypotheses’, ‘problems’, or ‘facts’ or
b. clauses that are tagged as ‘results’ or ‘implications’ that also contain external citations. and sentence *S*_2_ also contains either
  a. clauses that are goals or methods or
  b. results/implications with no external citations.
c. both *S*_1_ and *S*_2_ contain references to subfigures that are entirely disjoint (i.e., *S*_1_ refers to ‘Fig. 1C’ and *S*_2_ refers to ‘Fig. 1D, 1E and 1F’).
d. *S*_2_ is a section heading, indicating that the *S*_1_/*S*_2_ boundary marks a transition between sections.

Similarly, we repeated this process by scanning forward from the figure reference mention for the following conditions between consecutive sentences *S*_1_ and *S*_2_ indicating that *S*_1_ was the last sentence of the evidence fragment:

a. Sentence *S*_1_ contains only clauses that are tagged as as ‘results’ or ‘implications’ without citing external papers and Sentence *S*_2_ also contains only
  a. clauses that are tagged as ‘goals’, ‘methods’, ‘hypotheses’, ‘problems’, ‘facts’ or ‘methods’ or
  b. clauses that are tagged as ‘results’ or ‘implications’ with external citations present.

Conditions *b.* and *c.* headings were applied as before to detect the start of evidence fragments.

### 3.4 Applying the REACH event extraction tool

REACH is an event extraction engine for molecular signaling [5]. We applied REACH to INTACT open access papers and cross-referenced outputs to those linked to specific subfigures also referenced by INTACT data records. The only event type in REACH dealing with molecular interaction are ‘Complex Assembly’ events which we compared to data specified by INTACT data records to generate baseline event-extraction statistics.

### 3.5 Building the Molecular Interaction Evidence Fragment Corpus

We developed an OWL-based implementation of the existing BioC formulation [14], extended the SciDT pipeline system to export linked data conforming to that model. Also, we used the ‘Semantic Publishing and Referencing’ (SPAR) ontologies for bibliographic elements and references in both bioc and biopax linked data sets [15]. We used Paxtools [16] to convert INTACT PSI-MI2.5 data to BioPax (with a minor adaption to include figure references in the biopax representation of evidence).

## 4 Results

### 4.1 Discourse Tagging

In [12], Dasigi *et al.* evaluated 5-fold cross-validation Accuracies and F-Scores for SciDT based on a training set of 2,678 clauses over 263 paragraphs from results sections (Accuracy = 0.75, F-Score = 0.74). We extended this training data over all sections of the paper to yield 654 paragraphs with 6629 clauses. Of these, 253 paragraphs were from results sections yielding 2802 clauses.

### 4.2 Computing Figure Spans within Documents

Figure 3 illustrates the output of this procedure as a Gantt chart of the spans of subfigures over the clauses in a single paper’s results section. This shows how experiment references punctuate the argument of the paper with factual evidence. It also shows explicitly how a single paper in this domain is structured around a large number of small-scale experiments (23 in this case). We evaluated our methodology on a mixed set of manually annotated 10 open access papers (involving 190 figure references). This evaluation (of correctly identifying a figure reference for a given clause) gave macro average Precision = 0.66 *±* 0.02, Recall = 0.87 *±* 0.02 and F-score = 0.76 *±* 0.01.

**Figure 3.**
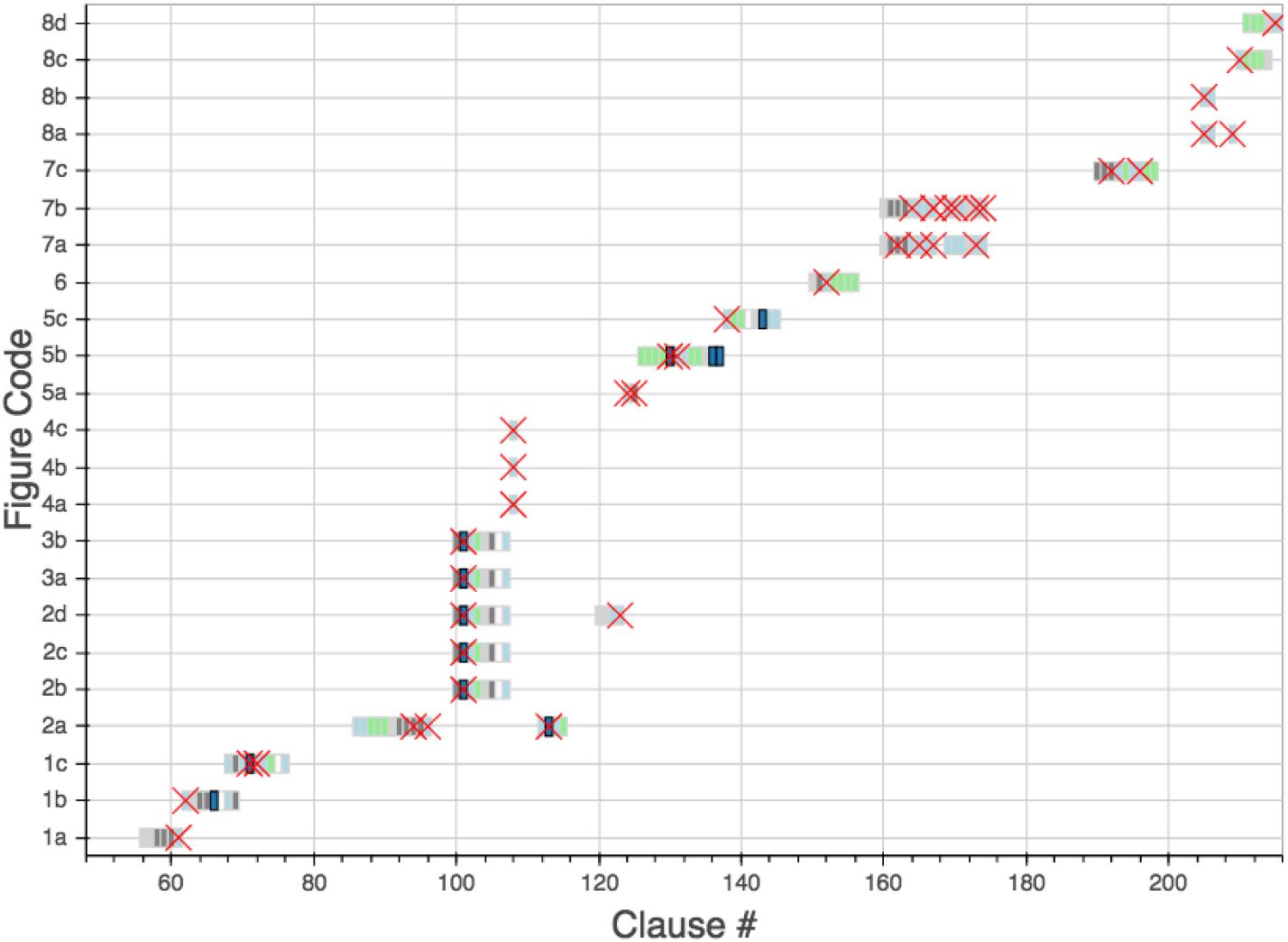
Gantt chart distribution of experimental spans for [13]. Red crosses show positions of subfigure references. Discourse type colors: ‘fact’/‘hypothesis’/‘problem’ = white; ‘goal’ = light gray; ‘method’ = gray; result = ‘light blue’; ‘implication’ = light green.

### 4.3 The Molecular Interaction Evidence Fragment Corpus

We have released all data associated with the study on FigShare [17]. The data consists of a compressed archive of individual files for papers’ evidence fragments and intact data records.

### 4.4 REACH System Output

We ran REACH over all available open source documents in INTACT. Of the 6320 INTACT records with associated figure references, we were able to identify a ‘Complex Assembly’ event within the sentences our system designated as associated with each event 2747 times (43.47% of records). The most precise measure of event extraction accuracy is based on matching the UNIPROT identifiers of any proteins described in the extracted REACH event to those of the INTACT data record. REACH was able to precisely reconstruct the INTACT data record to that level of accuracy in only 356 cases (5.6% of records). This provides a baseline measurement for future work.

## 5 Discussion

We have sought to instantiate a novel methodology for distant supervision in biomedical text mining and to provide the community access to a mid-sized text corpus for future use. Although our event extraction experiments showed poor performance, this provides a baseline for off-the-shelf tools that we expect to be able to improve upon straightforwardly. We would like to extend this to work with argumentation graphs where claims may be linked from other parts of papers [18,19]. Developing methods to automatically create such graphs *across* papers may provide powerful new ways of examining the literature.

Machine reading depends on the natural redundancy of any scientific narrative where common assertions are stated and restated in different ways across papers. On aggregate, these systems extract structured data from sentences that cite other work. This is problematic, since when evaluated for correctness, citation statements are often inaccurate [20]. More seriously, citations are both retained and reused within the literature even after the work that they are citing has been retracted [21]. Thus, a key, original focus of this work is to focus on the assertions that summarize the primary findings of a given paper rather than seek to use any and all available language to use for machine reading tasks.

## Acknowledgments.

This work was funded by DARPA Big Mechanism program under ARO contract W911NF-14-1-0436. We thank Anita de Waard, Mihai Surdeanu, Clay Morrison, and Hans Chalupsky for their contributions.

